# The Effect of Repeat Length on Marcal1-Dependent Single-Strand Annealing in *Drosophila*

**DOI:** 10.1101/2022.09.24.509346

**Authors:** Evan B. Dewey, Julie Korda Holsclaw, Kiyarash Saghaey, Mackenzie E. Wittmer, Jeff Sekelsky

## Abstract

Proper repair of DNA double strand breaks (DSBs) is essential to maintenance of genomic stability and avoidance of genetic disease. Organisms have many ways of repairing DSBs, including use of homologous sequences through homology-directed repair (HDR). While HDR repair is often error-free, in single-strand annealing (SSA) homologous repeats flanking a DSB are annealed to one another, leading to deletion of one repeat and the intervening sequences. Studies in yeast have shown a relationship between the length of the repeat and SSA efficacy. We sought to determine the effects of homology length on SSA in *Drosophila*, as *Drosophila* uses a different annealing enzyme (Marcal1) than yeast. Using an *in vivo* SSA assay, we show that 50 base pairs (bp) is insufficient to promote SSA and that 500-2000 bp is required for maximum efficiency. Loss of Marcal1 generally followed the same homology length trend as wild-type flies, with SSA frequencies reduced to about a third of wild-type frequencies regardless of homology length. Interestingly, we find a difference in SSA rates between 500 bp homologies that align to the annealing target either nearer or further from the DSB, a phenomenon that may be explained by Marcal1 dynamics. This study gives insights into Marcal1 function and provides important information to guide design of genome engineering strategies that use SSA to integrate linear DNA constructs into a chromosomal DSB.

## Introduction

Cellular DNA damage, particularly that caused by double-strand breaks (DSBs), must be properly cleared to maintain viability. DSBs arise from numerous intra- and extra-cellular sources, including external environmental chemical mutagens and ionizing radiation, as well as intracellular oxidative and replicative stress (Pfeiffer *et al*. 2000; Ciccia and Elledge 2010; Sage and Shikazono 2017). Left unrepaired, DSBs trigger apoptosis, and in improperly regulated cells aberrant repair can cause insertions, deletions, and translocations, leading to genomic instability and disease. Thus, it is critical for DSBs to be properly repaired to provide stability and avoid disease.

Cells have evolved robust mechanisms to repair DSBs accurately and avoid deleterious fates. Repair mechanisms are generally divided into two categories: non-homologous end joining (NHEJ) and homology-directed repair (HDR). While each strategy is effective in clearing DSB damage, each also has consequential tradeoffs. Canonical NHEJ does not require the homologous chromosome or sister chromatid as a template, instead requiring minimal processing of DSB ends prior to ligating them (Chang *et al*. 2017). This process is fast, simple, and efficient, but can frequently lead to small insertions and deletions.

HDR, if executed properly, provides benefits over NHEJ. Because HDR uses a homologous sequence as a template it can provide a largely error-free mechanism of repair. The cell commits to HDR over NHEJ by creating single-stranded DNA (ssDNA) tails with 3’-OH ends using specific exonucleases, a process called resection (reviewed in Cejka and Symington 2021). ssDNA tails are protected from degradation by replication protein A (RPA) before pursuing one of the many sub-pathways of HDR to complete repair. One such pathway is single-strand annealing (SSA), in which the ssDNA strands anneal to complementary sequence in one another and trim away any excess ssDNA flaps (Sugawara and Haber 1992; Rong and Golic 2003; Storici *et al*. 2006; Bhargava *et al*. 2016). As a result of the requirement for a large amount of complementary sequence, SSA occurs when a DSB is situated between direct repeats and is the one HDR pathway resulting in large deletions. In contrast to other HDR pathways, which generally have high fidelity, SSA necessarily generates a deletion.

Prior studies have identified proteins involved in SSA in fungi, invertebrates, and mammalian cell lines (reviewed in Bhargava *et al*. 2016; Vu *et al*. 2022). Studies in budding yeast found that loss of Rad52 or its paralog Rad59 resulted in near complete ablation of SSA (Sugawara and Haber 1992; Ivanov *et al*. 1996; Sugawara *et al*. 2000). Rad52 binds ssDNA with strong affinity and can displace the ssDNA binding protein RPA to facilitate annealing between complementary sequences, an activity likely important to promoting SSA (Shinohara and Ogawa 1998; Grimme *et al*. 2010; Ma *et al*. 2017). RAD52 has also been implicated in SSA in mammalian cells, though its loss doesn’t completely ablate SSA in these cells (Kelso *et al*. 2019).

While Rad52 functions in SSA in yeast and mammalian cells, it has been lost at several points in eukaryotic diversification, including in Dipteran insects (Sekelsky 2017). In *Drosophila*, Marcal1 is required for SSA (Korda Holsclaw 2017). Biochemical studies of Marcal1 demonstrate annealing activity promoted through a HepA-related protein (HARP) domain (Yusufzai and Kadonaga 2008; Kassavetis and Kadonaga 2014), and thus it seems likely that Marca1 has the same SSA role in flies as Rad52 in yeast. The human ortholog, SMARCAL1, has similar activities (Coleman *et al*. 2000; Ghosal *et al*. 2011), though a role in SSA has not been reported.

To gain insight into how Marcal1 promotes SSA through its annealing activity, we asked how changing the length of available complementarity would affect SSA efficiency in both wild-type and *Marcal1* mutant flies. This question is also of interest because SSA can be used for CRISPR/Cas9 based genome editing to incorporate large fragments into the *Drosophila* genome (Kanca *et al*. 2019). The first demonstration of this approach employed a plasmid with 100 nt of flanking homology to either side of two Cas9 genomic targets. The plasmid was cut *in vivo* to generate a linear fragment. While the authors were successful in integrating sequences with 100 nt of flanking homology, they speculated that increasing the homology would increase efficiency further without sacrificing the ease and low cost of synthesizing the constructs used.

Here we show that efficiency of SSA in *Drosophila* is indeed dependent on the length of homology available, with 50 bp being insufficient and more than 500 bp being required for optimal SSA. The amount of resection required to expose complementary sequences used in SSA do not affect SSA efficacy when large amounts of homology are present in wild type, though smaller amounts do appear to be more sensitive to resection distance. In flies lacking Marcal1, SSA success is significantly reduced compared to wild type regardless of length of homology present. However, even in the complete absence of Marcal1 there is substantial residual SSA, suggesting the existence of another annealing enzyme.

## Results

### SSA effectiveness is dependent on the length of homology present

To determine how length of homology affects efficiency of SSA in *Drosophila*, we built a modified *P*{*wIw*} assay (Rong and Golic 2003). In the original *P*{*wIw*}, two copies of the *mini-white* gene are inserted in tandem with an I-*Sce*I cut site in the middle (Figure 1A, top). The copy downstream (3’) of the I-*Sce*I cut site is functional, while the upstream (5’) copy is non-functional due to deletion of the promoter and part of the first exon. To perform the assay, I-*Sce*I is expressed via heat shock in the developing germline of males heterozygous for the insertion of *P*{*wIw*} on the third chromosome, and repair events recovered in progeny (Figure 1B-E). In previous studies, about 90% of progeny had cleavage and repair by SSA, suggesting that strand invasion and repair off the homologous chromosome is rare. Our modified assays retain this high I*-Sce*I cleavage efficacy (Figure S1).

**Figure 1.**
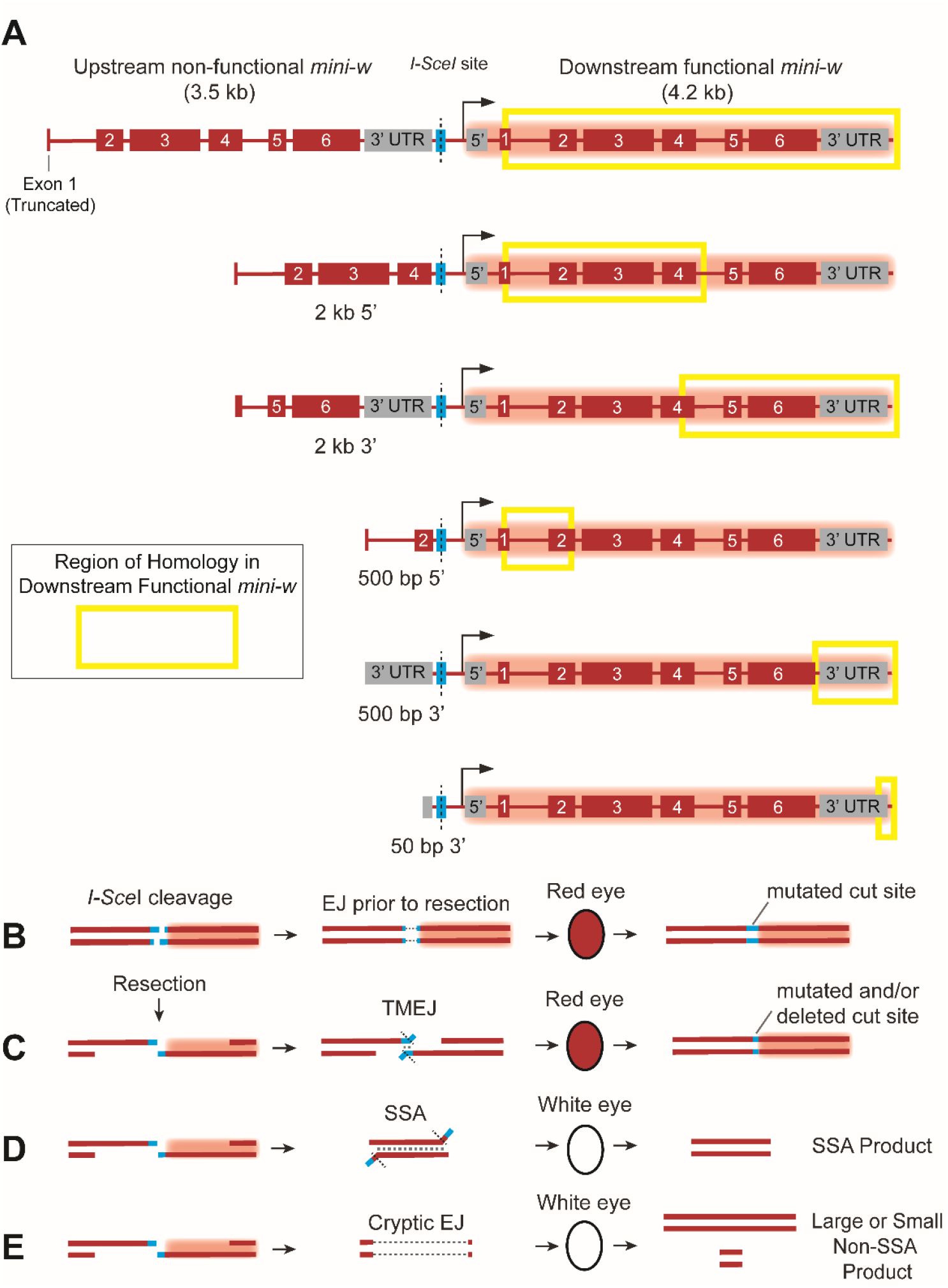
*P*{*wIw*} assay design and outcomes. (A) Homologies used in the *P*{*wIw*} assay. The original assay (Rong and Golic 2003) uses a non-functional *mini-white* gene with part of exon 1 deleted. This is upstream (5’) of the I-*Sce*I site. Our assay uses different upstream homology lengths corresponding to different regions of the functional, downstream (3’) *mini-w* gene (yellow boxes). (B-E) Outcomes of the assay. (B) Imprecise non-homologous end-joining (NHEJ/EJ) results in a mutated I*-Sce*I cut site and leads to a red eye in progeny. (C) Resection followed by DNA polymerase theta-mediated end joining (TMEJ) results in a mutated or deleted I-*Sce*I cut site, preventing further cutting and resulting in a red eye in progeny. (D) Resection followed by single-strand annealing (SSA) results in a distinct deletion for each homology class (SSA product) and white eyes in progeny. (E) Some deletions are larger than expected by canonical NHEJ or TMEJ; we refer to these are cryptic end joining since their origins have not been determined.

If the DSB is repaired by canonical NHEJ, resulting progeny will have red eyes (Figure 1B). The I-*Sce*I site in this case is often mutated due to the mutagenic nature of NHEJ, as non-mutated sites can be re-cut and re-repaired until a mutated, uncuttable outcome is produced (Figure 1B). Red eyes can also result form DNA polymerase theta-mediated end joining (TMEJ), a process that anneals microhomologies near the ends of resected DNA strands (Figure 1C) (Chan *et al*. 2010; Carvajal-Garcia *et al*. 2020). If resection on both sides of the break uncovers complementarity between the truncated upstream and full-length downstream *mini-white* genes, the repair product lacks a promoter and the first exon, resulting in progeny in white eyes among the progeny that inherit this SSA repair product (Figure 1D). White eyes may also result from larger deletions into the promoter or coding sequence of the downstream *mini-white*; such deletions can be detected by PCR and sequencing (Figure 1E).

To better elucidate *Marcal1*-based SSA mechanisms in *Drosophila*, we inserted *P*{*wIw*} constructs with upstream various non-functional *mini-white* fragments, creating five distinct versions (Figure 1A). We varied whether upstream homology was taken from the 5’ or 3’ portion of the *mini-white* gene, while keeping the distance from the downstream functional *mini-white* the same (Figure 1A, yellow boxes). With homologies taken from the 3’ part of the gene, more resection into the downstream *mini-white* gene is needed to expose complementary sequences, which we hypothesized could provide an additional constraint to effective SSA.

In wild-type flies, the percentage of white-eyed progeny (indicating likely SSA repair) was not significantly different between full length *P*{*wIw*} (3.5 kb of homology; 90.8%) and 2 kb of homology from either the 5’ (90.1%) or 3’ (89.0%) end of the gene (Figure 2A, B; Table 1). However, when homology from the downstream functional *mini-white* was reduced to 500 bp from either the 5’ or 3’ portion, there was a significant decrease in the number of white-eyed progeny compared to both full-length and 2 kb homologies (59.3% and 70.1%, respectively, *p* < 0.0001), indicating reduced SSA (Figure 2A, B; Table 1). Contrary to our hypothesis that 3’ homology might be additionally constraining on SSA due to the additional resection required, the opposite appeared to be true for the 500 bp homologies, with the 3’ 500 bp homology producing significantly more white-eyed progeny compared to the 5’ counterpart (Figure 2A, B; Table 1, p<0.001). Finally, when we reduced the amount of 5’ homology to 50 bp (from only the 3’ portion of the non-functional *mini-white* gene), we saw a significant decrease in white-eyed progeny compared to all other homology lengths (8.3%, *p* < 0.0001 for each comparison), indicating that 50 bp is below the homology threshold necessary for efficient SSA in this assay. While most of the white-eyed progeny in the *P*{*wIw*} assay result from SSA repair, white eyes may also result from non-SSA deletions into the functional *mini-white* gene (Figure 1E). To quantify this, we analyzed repair products in progeny. Because repair in the proliferating germline can result in multiple progeny from a single event, we assured independence of repair events by analyzing just one white-eyed offspring per vial, for a total of about 30 flies per experiment. Across all homology lengths except 50 bp (excluded as most vials did not have white-eyed progeny), SSA was the predominant form of repair (Figure 3A); for longer homologies (3.5 kb, 2 kb 5’, and 2 kb 3’), all white-eyed flies resulted from SSA (Figure 3B,C). For the 500 bp homologies we detected some non-SSA products, particularly in case of the 5’ homology (13.3% non-SSA; Figure 3D), suggesting SSA repair efficacy in *Drosophila* begins to diminish between 2 kb and 500 bp of homology. When we further multiply the white-eyed progeny for each condition by the percentage of SSA PCR products, we can obtain the true rate of SSA per condition (SSA Rate, Table 1). While SSA rates for homologies of 2 kb and above remain unchanged from the percentage of white-eyed progeny, as all white-eyed flies obtained yielded SSA products, we did see a modest decrease when comparing the percentage of white-eyed flies to the SSA rates for both 500 bp homologies, reflecting a more accurate rate of SSA for these conditions (Table 1). Last, for the 50 bp 3’ homology, we see that in vials with white-eyed flies, almost all repair products are non-SSA (90.9%, Figure S3), further indicating 50 bp is not enough homology to perform SSA efficiently in *Drosophila*.

**Table 1.** Comparisons of SSA Frequencies between wild-type and *Marcal1* mutant flies.

| Assay | Wild Type |  |  | *Marcal1* Mutant |  |  | Marcal1/<br>WT |
| --- | --- | --- | --- | --- | --- | --- | --- |
|  | White Eyes | SSA PCR | SSA Rate | White Eyes | SSA PCR | SSA Rate |  |
| 3.5 kb | 91 | 100 | 91 | 53 | 75 | 40 | 0.34 |
| 2 kb 5' | 90 | 100 | 90 | 42 | 53 | 22 | 0.24 |
| 2 kb 3' | 89 | 100 | 89 | 45 | 65 | 29 | 0.33 |
| 500 bp 5' | 59 | 87 | 51 | 28 | 80 | 22 | 0.43 |
| 500 bp 3' | 71 | 97 | 68 | 33 | 64 | 21 | 0.31 |

**Figure 2.**
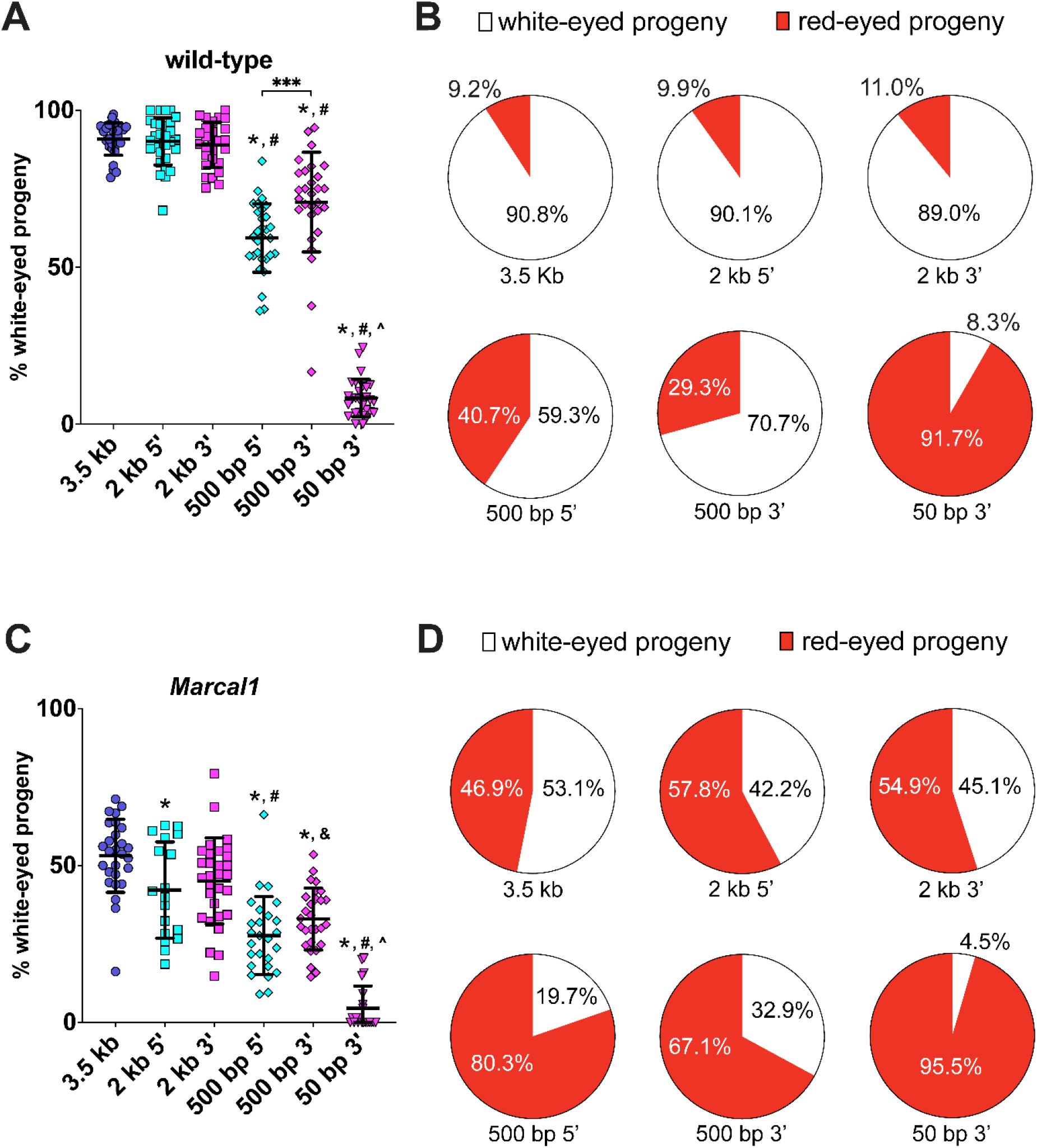
Length of homology affects SSA efficiency in wild-type and *Marcal1* mutant flies. (A) Percent of total progeny with white eyes in wild-type *P*{*wIw*} crosses. * *p* < 0.05 versus 3.5 kb homology, # *p* < 0.05 versus 2 kb homology (5’ and 3’), ^ *p* < 0.05 versus 500 bp homology (5’ and 3’), *** p<0.0001 between 500 bp 5’ and 500 bp 3’, based ANOVA with Tukey’s post hoc test. (B) Pie charts showing the white-eye vs. red-eye progeny for each wild-type assay. While the 3.5 kb and 2 kb homologies provide good templates for SSA (top row), as homology length is reduced, SSA efficiency diminishes (bottom row). (C) Percent of total progeny with white eyes in wild-type *P*{*wIw*} crosses. * *p* < 0.05 versus 3.5 kb homology, # *p* < 0.05 versus 2 kb homology (5’ and 3’), & *p* < 0.05 versus 2 kb 3’ homology, ^ *p* < 0.05 versus 500 bp homology (5’ and 3’), based on ANOVA with Tukey’s post hoc test. For 50 bp 3’, *n* = 1286 progeny; other *n* values are in Table 1. (D) Pie charts showing the white-eyed versus red-eyed progeny for each assay in *Marcal1* mutant flies.

**Figure 3.**
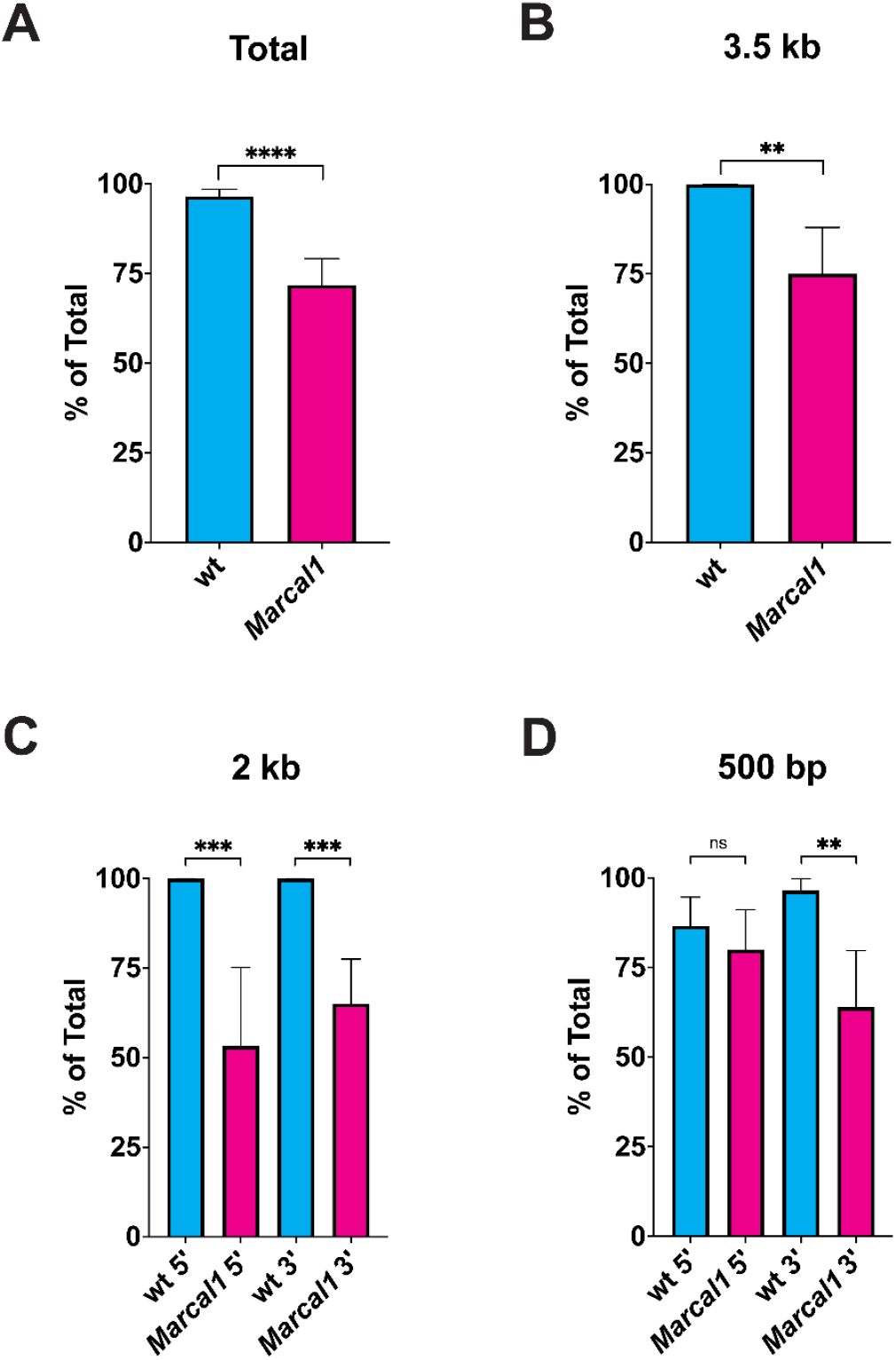
Molecular analysis of white-eyed progeny. In white-eyed progeny, the repaired region was amplified by PCR to determine whether region was repaired by SSA, producing a distinct product size or cryptic EJ, producing a larger or smaller product. Data from wild-type flies and *Marcal1* mutants were compared by Fisher’s exact test. (A) Summed data from all assays except 50 bp. n=144 for wt; n=117 for *Marcal1*; \*\*\*\**p*<0.0001. (B) 3.5 kb homology. *n*=29 for wild type, 24 for *Marcal1*; \*\**p*<0.01. (C) 2 kb 5’ and 3’ homologies. For 5’ *n*=30 for wild type, 15 for *Marcal1*; for 3’ *n*=26 for wild type, 43 for *Marcal1*; \*\*\**p*<0.001. (D) 500 bp 5’ and 3’ homologies. For 5’ *n*=30 for wild type, 25 for *Marcal1*; for 3’ *n*=29 for wild type, 25 for Marcal1; n.s., *p*=0.7165.

### Loss of Marcal1 reduces white-eyed progeny irrespective of homology length

We next asked whether differing lengths of homology affected the efficacy of SSA in a *Marcal1* mutant background. Our results also show a decrease of 41% for the assay with 3.5 kb homology when *Marcal1* is mutated (Figure 2C, D; Table 1). We speculated the effects of Marcal1 loss might become more severe as homology lengths were reduced; however, we found a 40-50% reduction in white-eyed progeny from wild type regardless of the amount of homology present (Figure 2C, D; Table 1). Overall, *Marcal1* mutants follow the same trend as wild type, with decreasing homology yielding decreasing amounts of white-eyed progeny (Figure 2C, D; Table 1).

### SSA repair events are less prevalent in Marcal1 white-eyed progeny

When initially examining the role of Marcal1 in SSA, Korda-Holsclaw and colleagues found that while the majority of white-eyed progeny from wild-type flies resulted from SSA repair, many of the white-eyed progeny resulting from *Marcal1* resulted from non-SSA deletions into the functional *mini-white* gene (Korda-Holsclaw, 2017). In agreement with this prior work, we found the percentage of white-eyed progeny that resulted from SSA repair was significantly different between wild type and *Marcal1* mutants (excluding 50 bp 3’ homology) across all homology amounts/lengths (96.5% and 71.8%, respectively; Figure 3A, Table 1, SSA Products).

When examining the percent of white-eyed flies with SSA products at differing lengths of homology, *Marcal1* mutant progeny do not follow the trend established in wild-type flies. While there is a decrease in the amount of SSA products when reducing homology from 3.5 kb to 2 kb 5’ and 3’ (Figure 3B,C), SSA products appear to remain the same or even increase when comparing the larger homologies (3.5 and 2 kb) to the 500 bp homologies (Figure 3B,C versus 3D, Table 1). When we multiply the percent of white-eyed progeny by the percent of SSA products to obtain the true SSA rate, we observe a decrease across all homologies (Table 1). Dividing the true SSA rates from *Marcal1* mutants by those from wild-type flies show that the decrease in SSA is stronger in Marcal1 flies when compared to the difference only in white-eyed progeny rates, varying from 44% to as low as 24% of wild type depending on condition (Marcal1/WT, Table 1). Together, these results indicate the effects of loss of Marcal1 on SSA are not strongly dependent on the length of homology.

## Discussion

### Homology amounts affect SSA

Here we demonstrate that homology length affects SSA in *Drosophila*. This agrees with previous studies in yeast (Sugawara and Haber 1992; Sugawara *et al*. 2000), and in human cell lines (Rothenberg *et al*. 2008; Kelso *et al*. 2019), although there are some differences across organisms in the threshold of homology required for SSA outcomes. Compared with our study, in which we begin to see a decrease in SSA efficacy in wild type between 2 kb and 500 bp, studies in yeast find a lower threshold, with this decrease starting between 905 and 415 bp (Sugawara and Haber 1992), or between 415 and 235 bp (Sugawara *et al*. 2000). This is less clear in human studies, as the variable lengths used are much smaller than in these and prior yeast experiments, topping out with homology lengths of 50 bp (Rothenberg *et al*. 2008) or 200 bp (Kelso *et al*. 2019). Whether higher amounts of homology yield better SSA rates in these contexts remains to be seen. Differences in the distances between repeats and their arrangements may also be a factor. Regardless, our work and others points to a homology threshold to efficiently carry out SSA. Factors involved in SSA may need a certain length to properly anneal repetitive sequence, with lower homology amounts not annealing as well as higher homology amounts. Stability gained through annealing could then allow for the accumulation of other SSA factors, such as endonucleases to cleave ssDNA flaps.

Why do organisms vary in their homology threshold amounts? This may be dependent on the primary annealing enzyme(s) involved in SSA; *Drosophila* Marcal1 may require longer stretches of complementarity than Rad52 in yeast and mammals. It is also possible that other SSA factors contribute to the differences.

We noted a significant difference in SSA repair between the 5’ 500 bp repeat and 3’ 500 bp repeat. GC content is similar, with 42% GC for 5’ 500 bp repeat compared to 40% for the 3’ 500 bp repeat. It is possible that the larger flap of ssDNA generated by annealing the 3’ 500 bp segments may be more favorable for cleavage (Figure 4A). The enzyme(s) that do this cleavage have not been identified, but Mei-9, the ortholog of the enzyme used in yeast, is not required for SSA in Drosophila (Wei and Rong 2007). Conversely, TMEJ may be favorable than SSA when the flap is shorter, as with the 5’ 500 bp segment. This could explain why more red-eyed flies are present in experiments with the 5’ 500 bp repeat, as a TMEJ mechanism would not be predicted to cause deletions into the downstream functional *mini-white* gene (see below, Carvajal-Garcia *et al*. 2020).

**Figure 4.**
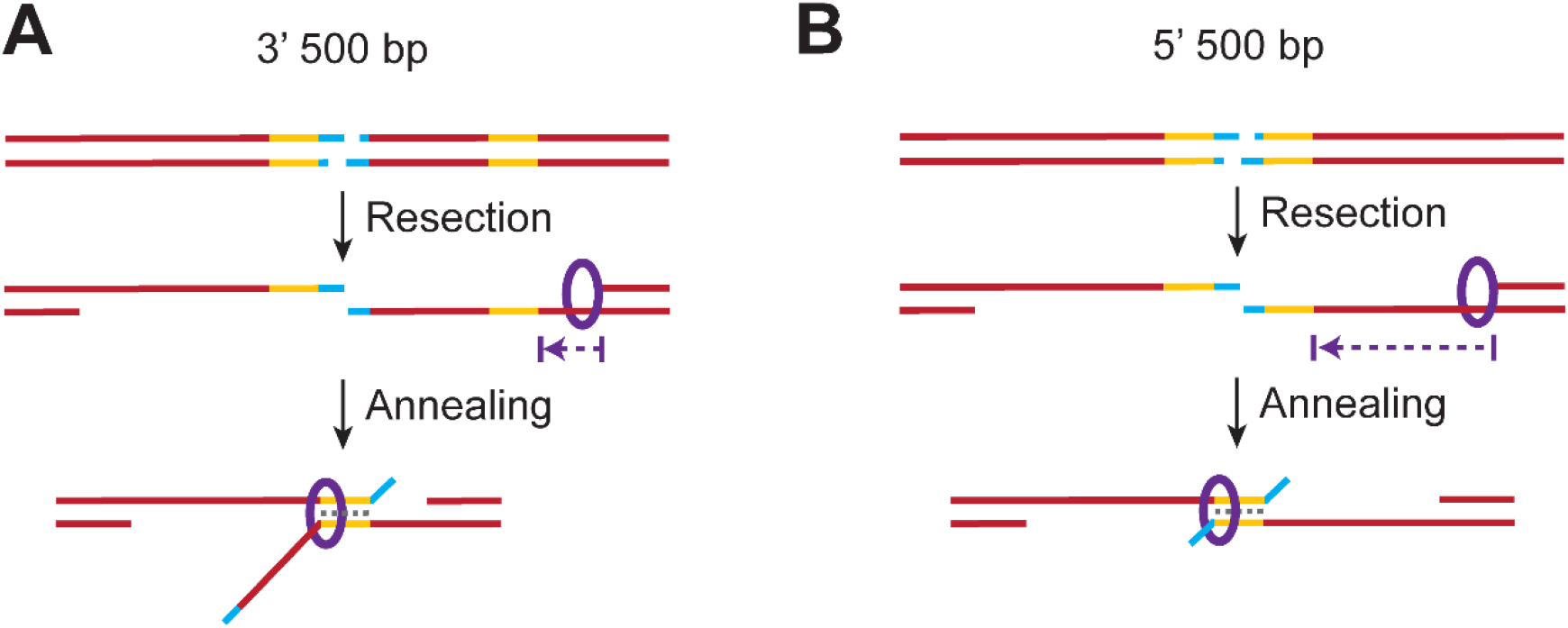
Possible mechanism for Marcal1 SSA based on differences between 5’ and 3’ 500 bp homologies. (A) Proposed Marcal1 mechanism for the 3’ 500 bp homology. Upon I-*Sce*I cutting and 5’ resection, Marcal1 (purple ring) localizes to the ssDNA-dsDNA interface created by resection. Marcal1 translocates along the ssDNA (indicated by purple arrow) until it finds a region of complementarity and promotes annealing. (B) Proposed Marcal1 mechanism for the 5’ 500 bp homology. Marcal1 localizes to the ssDNA-dsDNA interface created by resection and translocates towards the initial cut site. However, since the 5’ 500 bp homology of the functional *mini-white* gene is further from the ssDNA-dsDNA resection end point, Marcal1 must translocate a greater distance to reach homology and a higher probability of dissociation before reaching the region of complementarity. This may provide a longer window during which DNA polymerase theta can engage the ends to carry out TMEJ, resulting in a reduction in the SSA outcome.

Another possibility is that there is a preference to anneal repeats closer to the ssDNA-dsDNA interface created by resection. Based on high efficiency of the original *P*{*wIw*} assay, resection occurs at least 3.5 kb from the break site. By this mechanism, there would be more ssDNA between where the repeats anneal and the ssDNA-dsDNA interface for the 5’ 500 bp repeat, compared to less for the 3’ 500 bp repeat (Figure 4). This “interstitial” or “intervening” ssDNA could be more taxing for the cell to deal with, as there would be more of a ssDNA gap to fill following SSA with the 5’ 500 bp repeat versus the 3’ 500 bp repeat, making the 3’ 500 bp repeat more efficient.

A preference to anneal repeats closer to the ssDNA-dsDNA interface is an attractive model for *Drosophila* because of the binding characteristics of Marcal1. *In vitro* studies found that human SMARCAL1 and *Drosophila* Marcal1 bind to fork structures, which have an ssDNA-dsDNA interface, with higher affinity than to ssDNA and dsDNA (Yusufzai and Kadonaga 2008; Kassavetis and Kadonaga 2014). If Marcal1 is preferentially recruited to ssDNA-dsDNA interfaces, the start of the 3’ 500 bp ssDNA that would be annealed is closer to such an interface than for the 5’ 500 bp ssDNA (Figure 4 A, B). This would lower the ssDNA distance Marcal1 would need to traverse before annealing could be initiated, thereby favoring SSA. This preference would diminish with larger homologies used in our assays, as longer upstream homologies, even those matching the 5’ end of the downstream copy, decrease the distance between the ssDNA-dsDNA interface and annealing start site.

This mechanism would contrast Marcal1 with Rad52 function in several ways. First, each differs in how it binds to DNA. While Marcal1 has a preference to bind to ssDNA-dsDNA interfaces (Yusufzai and Kadonaga 2008; Kassavetis and Kadonaga 2014), yeast and human Rad52 has been shown to prefer ssDNA over ssDNA-dsDNA interfaces (Shinohara and Ogawa 1998; Navadgi *et al*. 2003; Rossi *et al*. 2021). Second, both proteins would facilitate annealing of homologous repeats differently. Rad52 forms oligomeric ring structures to bind ssDNA, with the rings then aggregating along the ssDNA (Shinohara and Ogawa 1998; Van Dyck *et al*. 1998; Stasiak *et al*. 2000; Ranatunga *et al*. 2001). The rings then interact with one another on opposing strands to facilitate annealing without the need to hydrolyze ATP (Ranatunga *et al*. 2001; Kagawa *et al*. 2008; Saotome *et al*. 2018). This contrasts with prior work in *Drosophila*, showing that SSA was affected by mutation of the Marcal1 ATP binding pocket, suggesting that Marcal1 needs to traverse DNA to facilitate annealing (Korda Holsclaw and Sekelsky 2017).

### Loss of Marcal1 decreases SSA repair rate versus wild type, regardless of homology length

Regardless of the amount of homology or the location of homology (5’ vs. 3’), mutation of *Marcal1* decreases SSA repair rates to between 56% and 76% versus wild type depending on homology (Figure 3, Table 1). This agrees with previous studies assaying *Marcal1* in SSA with full length repeats in the *P*{*wIw*} assay (Korda Holsclaw and Sekelsky 2017), which saw a roughly 50% decrease (93.0% white-eyed progeny for wild type versus 44.6% for *Marcal1*). In both the presence and absence of Marcal1, 50 bp remains insufficient to carry out SSA, and maximal SSA frequency is achieved between 500 and 2000 bp. This result suggests that *Drosophila* has other SSA annealing enzymes.

### Deletion frequency is increased in *Marcal1* mutants

Among the white-eyed progeny of *Marcal1* mutants, it was interesting to note that more non-SSA deletions were obtained. How might these deletions be facilitated? Canonical NHEJ can involve resection that requires the endonuclease Artemis (Biehs *et al*. 2017); however, *Drosophila* lacks an ortholog of this protein. While there is evidence for end processing in an NHEJ context in *Drosophila* (Bozas *et al*. 2009), there have yet to be any genes identified that might participate in this process. A 2007 screen by Wei and Rong (2007) did see some large deletions when NHEJ components (specifically *lig4*), were mutated in the *P*{*wIw*} assay. They suggested it could be that ends are more susceptible to degradation when NHEJ components are mutated.

TMEJ is also potential explanation for the increased deletions. However, the microhomology search extends only 15-20 nt from each end of the DSB, resulting in deletions of 34-40 bp (Carvajal-Garcia *et al*. 2020; Carvajal-Garcia *et al*. 2021). Given there are approximately 620 bp between the I-*Sce*I cut site and the functional white gene regardless of homology length in our assay, this would mean TMEJ deletions would likely not affect the functional *white* gene. It is therefore unlikely that TMEJ explains the large non-SSA deletions we observed among white-eyed progeny. It has been proposed that 3’ ssDNA can be lost by the 3’-5’ exonucleolytic function of DNA polymerase delta before polymerase theta can be engaged (Carvajal-Garcia *et al*. 2020; Carvajal-Garcia *et al*. 2021). This could explain some of the large deletions we observed, though this would be difficult to test because it would require a 3’-5’ exonuclease separation-of-function allele of DNA polymerase delta that retains the essential replication functions of the protein.

### SSA as a tool in CRISPR/Cas9 genome engineering in Drosophila

Kanca *et al*. (2019) showed that SSA can be used for CRISPR/Cas9 genome engineering in *Drosophila*. Although these authors achieved success with only 100 bp of homology on each side of the DSB, they suggested that this may not be optimal. Our work suggests that increasing this to more than 200 bp will enhance the success of integration, yielding more efficient gene edits per round of embryonic injections. Further studies can define the optimum conditions for CRISPR SSA integration that balance the needs of homology versus synthesis costs.

## Materials and Methods

### Cloning of *P*{*wIw*} constructs

Varying lengths of homology were made by reconstructing the *P*{*wIw*} construct used previously (Rong and Golic 2003; Wei and Rong 2007). We started with w+attB (previously deposited into AddGene, plasmid #30326), which has a fully functional *white* gene under control of a basal *Hsp70Bb* promoter (*Hsp70Bb::white*; referred to as *mini-white*). We added a full-length, non-functional *mini-white* gene with truncated exon 1 (3.5 kb) and a 3’ I-*Sce*I site upstream of the functional *mini-white* in the form of two gBlocks (Integrated DNA Technologies). These incorporated restriction enzyme sites such that cleavage and relegation would delete various segments of the upstream *mini-white* gene sequence. Deleting between *Hin*dIII sites left 2 kb 5’ homology; between two *Nhe*I sites left 2 kb 3’ homology; between two *Avr*II left 500 bp 5’ homology; between two *Age*I sites left 500 bp 3’ homology; and between two *Mlu*I sites left 50 bp 3’ homology. The two gBlocks were added to the w+attB vector by InFusion cloning (Clontech/Takara) into the *Hin*dIII site. Once the full length, non-functional 5’ copy was inserted, varying homologies described were created by cutting with a particular enzyme and ligating with T4 DNA ligase (New England Biolabs). All vectors were checked via Sanger sequencing to confirm proper amount/arrangement of upstream homology.

### Drosophila Stocks

Drosophila stocks were kept at 25°C on standard cornmeal medium (Archon Scientific). Mutant alleles used in this paper include *Marcal1*^*kh1*^ (Korda Holsclaw and Sekelsky 2017)and *Marcal1*^*del*^ (Baradaran-Heravi *et al*. 2012). Once all *P*{*wIw*} constructs were cloned, each was embryonically injected and integrated via phiC31 integration into *PBac*{*y+-attP-*3B}VK00031 (62E) (BestGene). Positive integrants were confirmed by expression of *mini-white* in a *y*^*2*^ *w*^*Δ*^ background. Integrants were balanced over *TM6B* and combined with *Marcal*^*kh1*^ mutants on chromosome *2*.

### P{wIw} Assay

The *P*{*wIw*} SSA repair assay was performed as described previously (Korda Holsclaw and Sekelsky 2017). Briefly, for each length/arrangement of homology, 4-5 *Marcal1*^*kh1*^/*CyO*; *P*{*wIw*}/*TM6B* virgin females were crossed to 3 *Marcal1*^*del*^/*CyO*; *Sb P*{*Hsp70Bb::I-SceI*}/*TM6B* males. One day later, first-instar larval progeny were heat-shocked at 37°C for 1 hour to induce expression of I-*Sce*I. Heat shock was repeated on the following day to ensure all first-instar larvae expressed I-*Sce*I. Larvae were then allowed to develop to adulthood, and *Marcal1*^*kh1*^/*Marcal1*^*del*^; *P*{*wIw*}/*Sb P*{*Hsp70Bb::I-SceI*} males were collected. These males were then crossed one at a time to 4-5 *y*^*2*^ *w*^*Δ*^ virgin females for three days and then discarded. Progeny from these crosses were then scored for red or white eyes. To ensure that continued cutting with I-*Sce*I did not occur throughout development, only *Sb*^*+*^ progeny were scored. Two red-eyed and two white-eyed flies were collected from each vial for analysis. DNA from each collected fly was extracted for analysis of the cut site (in red-eyed flies) and the repair products (in white-eyed flies).

To assess whether maternally deposited Marcal1 was affecting SSA in the zygotic embryos’ developing germlines, we additionally performed crosses with *Marcal1*^*kh1*^*/Marcal1*^*del*^; *P*{*wIw*}*/*TM6B females for four different homology lengths (3.5 kb, 2 kb 5’, 500 bp 5’ and 50 bp). We found no significant difference in white-eyed rates in males generated from this cross versus males generated from the initial cross with heterozygous mothers, indicating no detectable effect from maternally deposited Marcal1 on SSA (Supplemental Figure S2).

### PCR Analysis of red-eyed and white-eyed flies

To determine the cutting efficiency of I-*Sce*I for each length/arrangement of homology, the area surrounding and including the cut site was amplified by PCR from one red-eyed fly per vial with forward primer 5’-AGCGGATAACAATTTCACACAGG-3’ (M13R) and reverse primer 5’-AGCGGATAACAATTTCACACAGG-3’ (SSA_R) (see Supplemental Figure S1 for cutting efficiency). PCR products were subjected to cutting with I-*Sce*I (New England Biolabs) for 1 hour at 37°C then run on an agarose gel. If amplified DNA was uncut by I-*Sce*I, this indicates the *I-SceI* site was likely cut *in* vivo and then mutated/deleted through end-joining (NHEJ). If amplified DNA was cut by *I-SceI*, this indicates the *I-SceI* site was likely not cut *in* vivo or less likely joined without mutation/deletion by NHEJ. DNA successfully cut by *I-SceI* could not be distinguished between uncut and perfect NHEJ.

To determine the type of repair that occurred in white-eyed flies for length/arrangement homology, the area flanking non-functional 5’ and functional 3’ *mini*-*w* genes was amplified by PCR for one white-eyed fly per vial with forward primer 5’-GTTCGCTCAAATGGTTCCGA-3’ (pwIw_I-SceI_L) and reverse primer 5’-TCGCGATGTGTTCACTTTGT-3’ (pwIw_I-SceI_R). PCR products were then run on a gel and repair product type determined by size. Each length/arrangement of homology has a particular sized band predicted for an SSA repair product (meaning that SSA likely occurred *in vivo*), with larger and smaller bands classified as “non-SSA repair products,” meaning that these flies had a type of repair that caused a deletion into the functional white gene but not due to repair by SSA.

## Data Availability Statement

Plasmids, *Drosophila* stocks, and sequences are available upon request. The authors affirm that all data necessary for confirming the conclusions of the article are present within the article, figures, and table.

## Acknowledgements

We thank members of the Sekelsky lab for helpful comments on the manuscript. This work was supported by a grant from the National Institute of General Medical Sciences to JS under award 1R35GM118127. EBD was supported in part by a grants from the National Cancer Institute (T32 CA217824).

